# Linear Assembly of a Human Y Centromere

**DOI:** 10.1101/170373

**Authors:** Miten Jain, Hugh E. Olsen, Daniel J. Turner, David Stoddart, Kira V. Bulazel, Benedict Paten, David Haussler, Huntington F. Willard, Mark Akeson, Karen H. Miga

**Affiliations:** UC Santa Cruz Genomics Institute, University of California, Santa Cruz, CA, USA; Oxford Nanopore Technologies, Oxford, UK; Duke Institute for Genome Sciences and Policy, Duke University, Durham, North Carolina, USA; Geisinger National, Bethesda, Maryland, USA

## Abstract

The human genome reference sequence remains incomplete due to the challenge of assembling long tracts of near-identical tandem repeats-in centromeric regions. To address this, we have implemented a nanopore sequencing strategy to generate high quality reads that span hundreds of kilobases of highly repetitive DNAs. Here, we use this advance to produce a sequence assembly and characterization of the centromeric region of a human Y chromosome.

---

Centromeres are specialized loci that facilitate spindle attachment and ensure proper chromosome segregation during cell division. Normal human centromeric regions are defined by the enrichment of an AT-rich ~171-bp tandem repeat, known as alpha satellite DNA ^1^. The majority of alpha satellite DNAs are organized into higher order repeats (HORs), where chromosome-specific alpha satellite repeat units, or monomers, are reiterated as a single repeat structure hundreds or thousands of times with high (>99%) sequence conservation to form extensive arrays ^2^. The sequence composition of individual HOR structures and the extent of repeat variation within the context of each chromosome-assigned HOR array are important to establish kinetochore assembly and ensure centromere identity ^3-5^. Yet, despite the functional significance of the genomic organization and structure, sequences within human centromeric regions remain absent from even the most complete chromosome assemblies. To date, no sequencing technology (e.g. SMRT sequencing or synthetic long reads technologies) or collection of sequencing technologies has been capable of assembling through centromeric regions due to the requirements for extremely high quality, long reads to confidently traverse low-copy sequence variants within a given array. To this end, we have implemented a nanopore long read sequencing strategy to generate high-quality reads capable of spanning hundreds of kilobases of highly repetitive DNAs (Supplementary Fig 1). We have focused on the haploid satellite array that spans the Y centromere (DYZ3) as it is particularly suitable for assembly due to its tractable array size, well-characterized HOR structure and previous physical mapping data^6-8^

We employed a transposase-based method (defined here as our ‘Longboard Strategy’) to generate high-read coverage of full-length bacterial artificial chromosome (BAC) DNA with nanopore sequencing (MinION sequencing device, Oxford Nanopore Technologies). This method is designed to linearize the circular BAC with a single cut-site, followed by addition of the necessary sequencing adaptors (Fig. 1a). The BAC DNA is then read in its entirety through the pore, resulting in complete, end-to-end read coverage of the BAC insert sequence. Plots of read length versus megabase yield revealed enrichment for full length BAC DNA sequences (Fig. 1b; Supplementary Fig. 2). In total, we generated over 3500 full-length “1D” reads (i.e. sequencing one strand of the DNA) that span the entirety of 10 BACs (two control BACs from Xq24 and Yp11.2; and eight BACs that mapped to the DYZ3 locus ^9^) with MinION sequencing (Supplementary Table 1).

**Figure 1:**
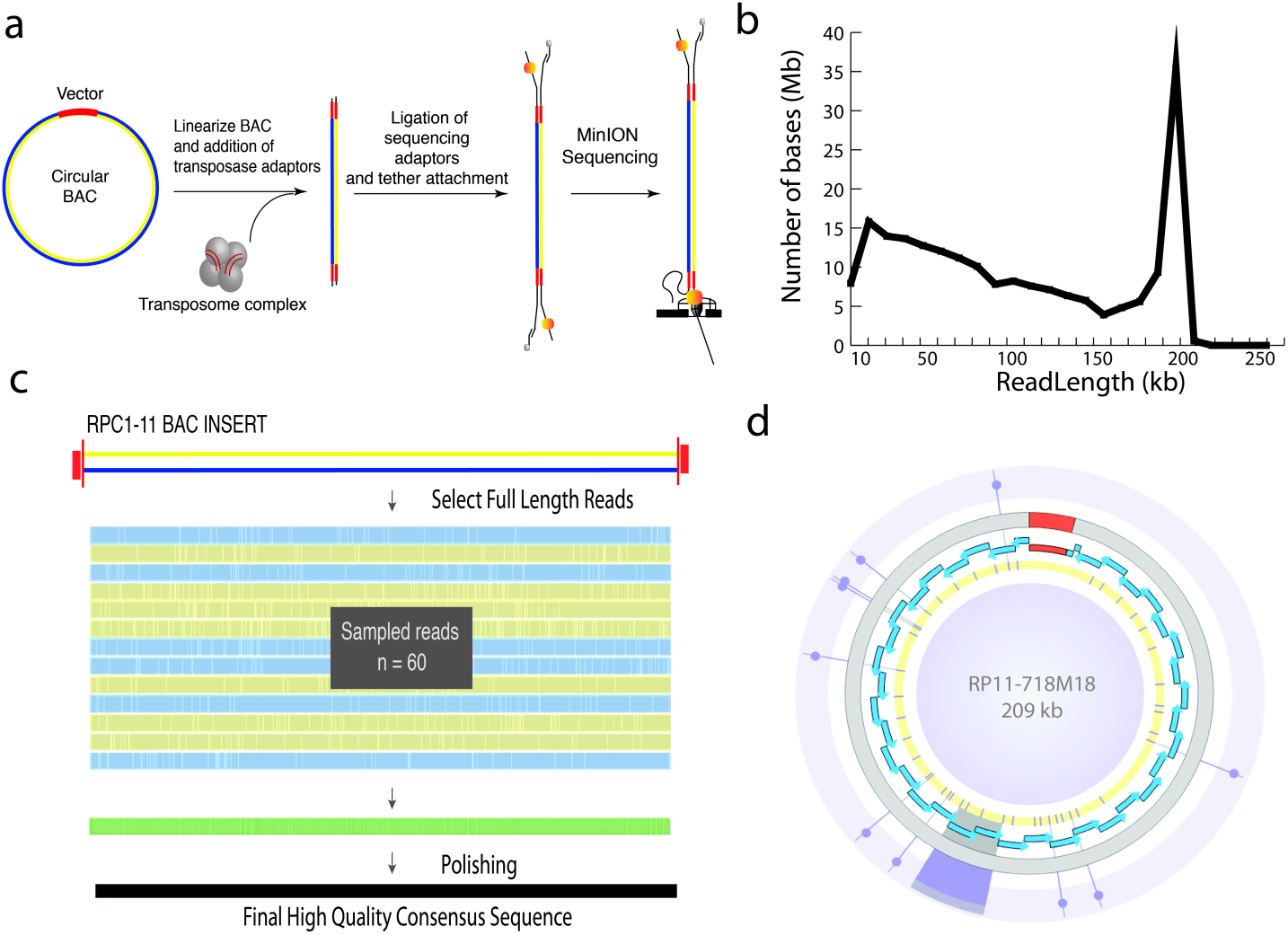
BAC-based Longboard nanopore sequencing strategy on the MinION. (a) Optimized strategy to cut each circular BAC once with transposase, resulting in a linear and complete DNA fragment of the BAC for nanopore sequencing. (b) Yield plots of BAC DNA (RP11-648J18) provide enrichment supporting BAC lengths. (c) High quality BAC consensus sequences were generated by multiple alignment of 60 full length 1D reads (shown as blue and yellow for both orientations) sampled at random with 10 iterations, followed by polishing steps (green) with the entire nanopore long read data and Illumina data. (d) Representation ^20^ of the polished RP11-718M18 BAC consensus sequence. Blue arrowheads indicate the position and orientation of HORs. Purple tiles in yellow background mark the position of the Illumina-validated variants. Additional purple highlight extending from select illumina-validated variants are used to identify single nucleotide sequence variants and mark the site of the DYZ3 repeat structural variants (6 kb) in tandem.

Correct assembly across the centromeric locus requires overlap among a few sequence variants, thus placing great importance on the accuracy of base-calls. Individual reads (MinION R9.4 chemistry, Albacore v1.1.1) provide inadequate sequence identity (i.e. median alignment identity of 84.8% for control BAC, RP11-482A22 reads) to ensure proper repeat assembly ^10^. To improve overall base quality, we derived a consensus from multiple alignments of 1D reads (60 randomly sampled full-length reads, with 10 iterations) that span the full insert length for each BAC (Fig. 1c). To polish sequence base quality, full-length nanopore reads were realigned to each BAC derived consensus (99.2% observed for control BAC, RP11-482A22; and an observed range of 99.4 - 99.8% for vector sequences in DYZ3-containing BACs). To provide a truth set of array sequence variants and to evaluate any inherent nanopore sequence biases, we performed Illumina BAC resequencing (Online Methods). As a result, we report nine BAC polished sequences, (e.g. 209 kb for RP11-718M18, Fig. 1d) to guide the ordered assembly of BACs from p-arm to q-arm, spanning an entire Y centromere.

We ordered the DYZ3-containing BACs using 16 Illumina-validated HOR variants, resulting in 365 kb of assembled alpha satellite DNA (Fig. 2a; Supplementary Data 1). The centromeric locus is dominated by a 301 kb array that is comprised entirely of the DYZ3 higher-order repeat (HOR), with a 5.8 kb consensus sequence, repeated in a head-to-tail orientation without repeat inversions or transposable element interruptions ^6,11,12^. The assembled length of the RP11 DYZ3 array is consistent with estimates for 96 individuals from the same Y haplogroup (R1b) (Supplementary Fig. 3; mean: 315 kb; median: 350 kb) ^13,14^. This finding is in general agreement with pulse-field gel electrophoresis (PFGE) DYZ3 size estimates from previous physical maps, and using a Y-haplogroup matched cell line (Supplementary Fig. 4).

**Figure 2:**
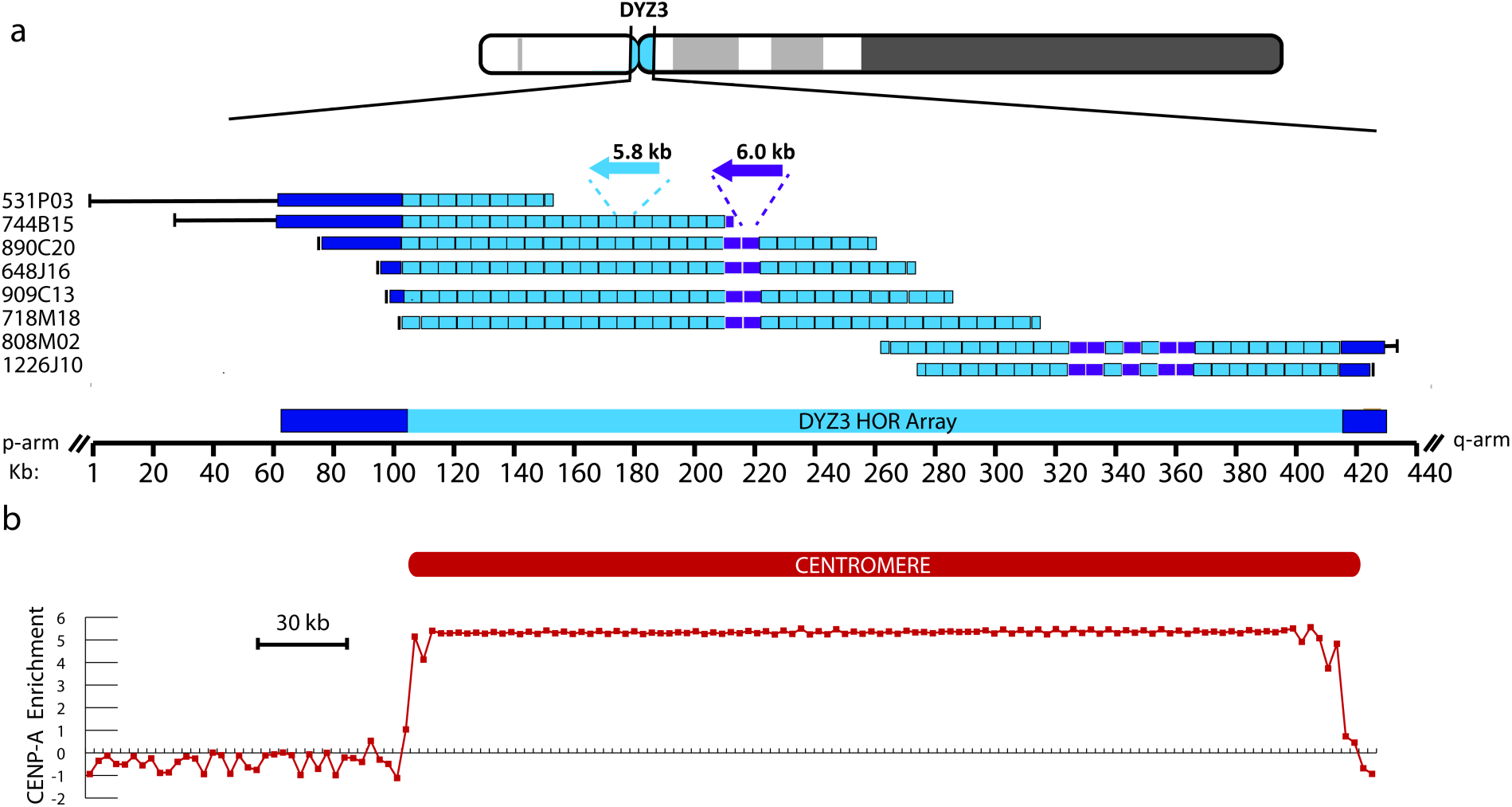
Linear assembly of the RP11 Y centromere. (a) Ordering of nine DYZ3-containing BACs spanning from proximal p-arm to proximal q-arm. The majority of the centromeric locus is defined by the DYZ3 conical 5.8 kb higher-order repeat (HOR) (light blue). Highly divergent monomeric alpha satellite is indicated in dark blue. HOR variants (6.0 kb) indicated in purple. (b) The genomic location of the functional Y centromere is defined by the enrichment of centromere protein A (CENP-A), where enrichment (~5-6x) is attributed predominantly to the DYZ3 HOR array.

Pairwise comparisons among the 52 HORs in the assembled DYZ3 array reveal limited sequence divergence between copies (mean 99.7% pairwise identity). In agreement with previous assessment of sequence variation within the DYZ3 array ^6^, we detected instances of a 6.0 kb HOR structural variant and provide evidence for seven copies within the RP11 DYZ3 array that are present in two clusters separated by 110 kb, as roughly predicted by previous restriction map estimates ^8^. Sequence characterization of the DYZ3 array revealed nine HOR haplotypes, defined by linkage between variant bases that are frequent in the array (Supplementary Fig. 5). These HOR-haplotypes are organized into three local blocks that are enriched for distinct haplotype groups, consistent with previous demonstrations of short-range homogenization of satellite DNA sequence variants ^6,15,16^.

Functional centromeres are defined by the presence of inner centromere proteins that epigenetically mark the site of kinetochore assembly ^17–19^. To define the genomic position of the functional centromere on the Y chromosome we performed an initial study of the enrichment profiles of inner kinetochore centromere protein A (CENP-A), a histone H3 variant that replaces histone H3 in centromeric nucleosomes, using a Y-haplogroup matched cell line that offers a similar DYZ3 array sequence (Fig. 2b, Supplementary Data 2) ^5,14,19^ In doing so, we find that CENP-A enrichment is predominantly restricted to the canonical DYZ3 HOR array, although we do identify reduced centromere protein enrichment extending up to 20 kb into flanking divergent alpha satellite on both the p-arm and q-arm side. Thus, we have provided a complete genomic definition of a human centromere, critical to advance sequence-based studies of centromere identity and function.

In conclusion, we have implemented a long read strategy to advance sequence characterization of tandemly repeated satellite DNAs. Despite their repetitive content, our analysis provides the necessary directed genomic approach to map, sequence and assemble centromere regions. In doing so, we report the array repeat organization and structure of a human centromere on chromosome Y. Previous modeled satellite arrays ^14^ are based on incomplete and gapped maps, and do not present complete assembly data across the full array. Only with a complete assembly, aided by longer reads, can one robustly measure the precise number of repeats in an array and resolve the order, orientation, and density of both repeat-length variants across the full extent of the array, critical to advance studies of centromere evolution and function. We expect that this work will be applicable to ongoing efforts to complete the human genome.

## Data Availability

Sequence data that support the findings of this study have been deposited in GenBank with reference to BioProject ID PRJNA397218, and SRA accession codes: - SRR5902337-SRR5902355. BAC consensus sequences and RP11-CENY array assembly GenBank Accessions: MF741337-47.

## Acknowledgements

This work was supported by grants to M.A from NHGRI [HG007827] and D.H., B.P. [DT06172015] from the Keck Foundation.

## Contributions

K.M and H.W. conceived the project. K.M., M.J., D.T., D.S., H.O. and M.A. designed the experiments; M.J. and H.O. were involved with BAC sample preparation; M.J. and H.O. performed MinION sequencing and base-calling; M.J and K.M analyzed the BAC sequencing data and validation analyses; K.M. performed the pulse-field gel electrophoresis array length estimates; K.B. contributed FISH analysis; K.M., M.J. and H.O. contributed to analysis and figure generation; M.A., D.T., D.S., H.W., B.P., and D.H. provided technical advice; all authors contributed to the writing, editing and completion of the manuscript.

## Online Methods

### I. Longboard MinION Protocol

#### BAC DNA Preparation and Validation

Bacterial artificial chromosomes (BACs) clones used in this study were obtained from BACPAC RPC1-11 library, Children's Hospital Oakland Research Institute in Oakland, California, USA (http://www.chori.org/bacpac/). BACs that span the human Y centromere: RP11-108I14, RP11-1226J10, RP11-808M02, RP11-531P03, RP11-909C13, RP11-890C20, RP11-744B15, RP11-648J16, RP11-718M18, and RP11-482A22, were determined based on previous hybridization with DYZ3-specific probes, and confirmed by PCR with STSs sY715 and sY78 ^9^. Notably, DYZ3 sequences, unlike shorter satellite DNAs ^21,22^, have been observed to be stable and cloned without bias ^5,23^. The RP11-482A22 BAC was selected as our control since it had previously been characterized by nanopore long-read sequencing ^24^ and presented ~134 kb of assembled, unique sequence present in the GRCh38 reference assembly to evaluate our alignment and polishing strategy. BAC DNA was prepared using the QIAGEN Large-Construct Kit (Cat No./ID: 12462). To ensure removal of the *E.coli* genome, it was important to include an exonuclease incubation step at 37°C for 1 hour, as provided within the QIAGEN Large-Construct Kit. BAC DNAs were hydrated in TE buffer. BAC Insert length estimates were determined by pulsed-field gel electrophoresis (PFGE) (data not shown).

#### Transposase-mediated 1D long reads

MinIONs can process long fragments, as has been previously documented ^24^. While these long reads demonstrate the processivity of nanopore sequencing, they are also few in numbers. To systematically enrich for the number of long reads per MinION sequencing run, we developed a strategy that uses the ONT Rapid Sequencing Kit (RAD002). We performed a titration between the transposase from this Kit (RAD002) and circular BAC DNA. This was done to achieve conditions that would optimize the probability of individual circular BAC fragments being cut by the transposase only once. To this end, we diluted the ‘live’ transposase from the RAD002 kit with the ‘dead’ transposase provided by ONT. For pulsed-field gel electrophoresis (PFGE) based tests, we used 1 μl of ‘live’ transposase and 1.5 μl of ‘dead’ transposase per 200 ng of DNA in a 10 μl reaction volume. This reaction mix was then incubated at 30°C for 1 minute and 75°C for 1 minute, followed by PFGE. Our PFGE tests used a 1% high-melting agarose gels and were run with standard 180° FIGE conditions for 3.5 hours. An example PFGE gel is shown in Supplementary Fig 6.

For MinION sequencing library preparation, we used 1.5 μl of ‘live’ transposase and 1 μl of ‘dead’ transposase (supplied by ONT) per 1 μg of DNA in a 10 μl reaction volume. Briefly, this reaction mix was then incubated at 30°C for 1 minute and 75°C for 1 minute. We then added 1 μl of the sequencing adapter and 1 μl of Blunt/TA Ligase Master Mix (New England Biolabs) and incubated the reaction for 5 minutes. This was the adapted BAC DNA library for the MinION. R9.4 SpotON flow cells were primed using the ONT recommended protocol. We prepared 1 ml of priming buffer with a 500 μl running buffer (RBF) and 500 μl water. Flow cells were primed with 800 μl priming buffer via the side loading port. We waited for 5 minutes to ensure initial buffering before loading the remaining 200 μl of priming buffer via the side loading port but with the SpotON open. We next added 35 μl RBF and 28 μl water to the 12 μl library for a total volume of 75 μΚ We loaded this library on the flow cell via the SpotON port and proceeded to start a 48 hour MinION run.

When a nanopore run is underway, the amplifiers controlling individual pores can alter voltage to get rid of unadapted molecules which can otherwise block the pore. With R9.4 chemistry, ONT introduced global flicking that reversed the potential every 10 minutes by default to clear all nanopores of all molecules. At 450 bps speed, a 200 kb BAC would take around 7.5 minutes to process. To ensure sufficient time for capturing BAC molecules on the MinION, we changed the global flicking time period to 30 minutes. This is no longer the case with an update to ONT’s MinKNOW software, and on the later BAC sequencing runs we did not change any parameters. We acknowledge that generating long (>100kb) reads presents challenges given the dynamics of HMW DNA for ligation, chemistry updates, and delivery of free ends to the pore reducing the effective yield. We found that high quality and quantity of starting material (i.e. our strategy is designed for 1ug of starting material that does not show signs of DNA shearing and/or degradation when evaluated by PFGE) and reduction of smaller DNA fragments were necessary for the Longboard Strategy.

### II. Protocol to improve long read sequence by consensus and polishing

#### Brief overview

BAC-based assembly across the DYZ3 locus requires overlap among a few informative sequence variants, thus placing great importance on the accuracy of base-calls. Therefore, we employed the following strategy to improve overall base quality. First, we derived a consensus from multiple alignments of 1D reads that span the full insert length for each BAC. Further, polishing steps were performed using re-alignment of all full-length nanopore reads for each BAC. As a result, each BAC sequencing project resulted in a single polished, BAC consensus sequence. To validate single copy variants, useful in a overlap-layout-assembly strategy, we included Illumina datasets for each BAC. Illumina data was not used to correct or validate variants observed multiple times within a given BAC sequence due to the reduced mapping quality.

#### A. MinION Base calling

All of the BAC runs were initially base-called using Metrichor, ONT’s cloud basecaller. Metrichor classified reads as pass or fail using a Q-value threshold. We selected the full-length BAC reads from the pass reads. We later rebase-called all of the BAC runs using Albacore 1.1.1 which included significant improvements on homopolymer calls. This version of Albacore did not contain a pass/fail cutoff. We reperformed the informatics using Albacore base-calls for full-length reads selected from the pass Metrichor base-calls. We selected BAC full-length reads as determined by observed enrichment in our yield plots (shown in Supplementary Fig 7 the read vs read length plots converted to yield plots to identify BAC length min-max selection thresholds).

Full-length reads used in this study were determined to contain at least 3 kb of vector sequence, as determined by BLASR ^25^ (*-sdpTupleSize 8 ‐bestn 1 ‐nproc 8 ‐m 0*) alignment with the pBACe3.6 vector (GenBank Accession: U80929.2). Reads were converted to the forward strand. Reads were reoriented relative to a fixed 3 kb vector sequence, aligning the transition from vector to insert.

#### B. Derive BAC Consensus Sequence

Reoriented reads were sampled at random (blasr_output.py). Multiple sequence alignment (MSA) was performed using kalign ^26^. We determined empirically that sampling greater than 60 reads provided limited benefit to consensus base quality (as shown in Supplementary Fig. 8). We computed the consensus from the MSA whereas the most prevalent base at each position was called. Gaps were only considered in the consensus if the second most frequent nucleotide at that position was present in less than 10 reads. We performed random sampling followed by MSA iteratively 10x, resulting in a panel of 10 consensus sequences, observed to provide a ~1% boost in consensus sequence identity (as shown in Supplementary Fig. 8). To improve the final consensus sequence, we next performed a final MSA on the collection of 10 consensus sequences derived from sampling.

#### C. Polish BAC Consensus sequence

Consensus sequence polishing was performed by aligning full-length 1D nanopore reads for each BAC to the consensus (BLASR^25^, *-sdpTupleSize 8 ‐bestn 1 ‐nproc 8 ‐m 0*). We used pysamstats (https://github.com/alimanfoo/pysamstats) to identify read support for each base call. We determined the average base coverage for each back, and filtered those bases that had low-coverage support (defined as having less than half of the average base coverage). Bases were lower-case masked if they were supported by sufficient sequence coverage, yet had less than 50% support for a given base call in the reads aligned.

#### D. Variant Validation

We performed Illumina re-sequencing (Miseq V3 600bp; 2 x 300 bp) for all nine DYZ3-containing BACs to validate single copy DYZ3 HOR variants in the nanopore consensus sequence. Inherent sequence bias is expected in nanopore sequencing ^24^, therefore we first used the Illumina matched datasets to evaluate the extent and type of sequence bias in our initial read sets, and our final polished consensus sequence. Changes in ionic current as individual DNA strands are read through the nanopore are each associated with a unique five-nucleotide k-mer. Therefore in an effort to detect inherent sequence errors due to nanopore sensing, we compared counts of 5-mers. Alignment of full length HORs within each polished BAC sequence to the canonical DYZ3 repeat demonstrated that these sequences are nearly-identical, where in RP11-718M18 we detected 1449 variant positions (42% mismatches, 27% deletions, and 31% insertions) across 202,582 bp of repeats (99.5% identity). Although the 5-mer frequency profiles between the two datasets were largely concordant (as shown in Supplementary Fig 9), we found that poly(dA) and poly(dT) homopolymers were overrepresented in our initial nanopore read datasets, a finding that is consistent with genome-wide observations. These poly(dA) and poly(dT) over-representations were reduced in our quality corrected consensus sequences especially for 6-mers and 7-mers.

#### D1. K-mer method

Using a k-mer strategy (where k=21 bp), we identified exact matches between the Illumina and each BAC consensus sequence. Illumina read data and the BAC polished consensus sequences were reformatted into respective k-mer library (where k=21 bp, with 1 bp slide using Jellyfish v2 software ^27^) in forward and reverse orientation. K-mers that matched the pBACe3.6 sequence exactly were labeled as ‘vector’. K-mers that matched the DYZ3 consensus sequence exactly ^14^ were labeled as ‘ceny’. We first demonstrated that the labeled k-mers were useful in predicting copy number. Initially, we showed how the ceny k-mer frequency in the BACs predicted the DYZ3 copy number, relative to the number observed in our nanopore consensus (as shown in figure below, panel a). DYZ3 copy number in each consensus sequence derived from nanopore reads was determined using HMMER3 ^28^ (v3.1b2) with a profile constructed from the DYZ3 reference repeat. By plotting the distribution of vector k-mer counts (shown in figure below, panel b for RP11-718M18), we observe a range of expected k-mer counts for a single copy sites. DYZ3 repeat variants (single copy satVARs) were determined as k-mers that (1) did not identified to have an exact match with either the vector or DYZ3 reference repeat, (2) spanned a single DYZ3 assigned variant in reference polished consensus sequence (i.e. that particular k-mer was observed only once in the reference), (3) and had a k-mer depth profile in the range of the corresponding BAC vector k-mer distribution. As a final conservative measure, satVARs used in overlap-layout-consensus assembly were supported by 2 or more overlapping illumina k-mers (shown in Supplementary Figure 10). To test if it was possible to predict a single copy DYZ3 repeat variant by chance, or by error introduced in the Illumina read sequences, we ran 1000 simulated trials using our RP11-718M18 Illumina data. Here, we randomly introduced a single variant into the polished RP11-718M18 DYZ3 array (false positive). We generated 1000 simulated sequences, each containing a single randomly introduced single copy variant. Next, we queried if the 21-mer spanning the introduced variant was (a) found in the corresponding Illumina dataset and (b) if so, monitored the coverage. Ultimately, none of the simulated false positive variants (21-mer) met our criteria of a true variant. That is, although the simulated variants were identified in our Illumina data, they had insufficient sequence coverage to be included in our study. Greater than 95% of the introduced false variants had less or equal to 100x coverage, with only one variant observed to have the maximum value of 300x. True variants were determined using this dataset with values between 1100-1600x as observed in our vector distribution.

#### D2. Alignment method

In addition to our k-mer based strategy, we also employed a short read alignment strategy to validate single copy variants in our polished consensus sequence. Illumina merged reads (PEAR, standard parameters ^29^) were mapped to the RP11 Y-assembled sequence using BWA-MEM. BWA-MEM is a component of the BWA package and was chosen because of its speed and ubiquitous use in sequence mapping and analysis pipelines. Aside from the difficulties of mapping the ultra-long reads unique to this work, any other mapper could be used instead. This involves mapping Illumina data to each BAC consensus sequence. After filtering those alignments with mapping quality less than 20, single nucleotide DYZ3 variants (i.e. a variant that is observed uniquely, or once in a DYZ3 HOR in a given BAC) are considered “validated” if they have (a) support of at least 80% of the reads and (b) have sequence coverage within the read depth distribution observed in the single copy vector sequence for each BAC dataset.

To explore illumina sequence coverage necessary for our consensus polishing strategy we initially investigated a range (20-100x) of simulated sequence coverage relative to a 73 kb control region (hg38 chrY:10137141-10210167) within the RP11-531P03 BAC data. Simulated paired read data using the ART illumina simulator software (Huang, 2011) was specified for the MiSeq sequencing system (MiSeq v3 (250bp), or ‘MSv3’), with a mean size of 400 bp DNA fragments for paired-end simulations. Using our polishing protocol, where: (1) reads are filtered by mapping quality score (ie. at least a score of 20: that the probability of correctly mapping is log10 of 0.01 * ‐10, or 0.99). (2) base frequency was next determined for each position using pysamstats, and (3) a final, polished consensus was determined by taking the base call at any given position that is represented by sufficient coverage (at least half of the determined average across the entire BAC) and is supported by a percentage of illumina reads mapped to that location (in our study, we require at least 80%). If we require at least 80% of mapped reads to support a given base call, we determine that 30x coverage is sufficient to reach 99% sequence identity (or the same as our observed identity using our entire Illumina read dataset, indicated as a grey dotted line in Supplementary Fig 11). If we require at least 90% of mapped reads to support a particular variant it is necessary to increase coverage to 70x to reach an equivalent polished percent identity.

To evaluate our mapping strategy we performed a basic simulation using a artificially generated array of 10 identical DYZ3 (5.7 kb) repeats. We then randomly introduced a single base change resulting in a new sequence with 9 identical DYZ3 repeats and one repeat distinguished by a single nucleotide change (as illustrated in Supplementary Fig 12). We first demonstrate that we are able to confidently detect the single variant by simulating reads from the reference sequence containing the introduced variant of varying coverage and Illumina substitution error rate. Additionally, we investigated whether we would detect the variant as an artifact due to illumina read errors. To test this, we next simulated Illumina reads from a DYZ3 reference array that did not contain the introduce variant (i.e. 10 exact copies of the DYZ3 repeat). We performed this simulation 100x, thus creating 100x reference arrays each with a randomly placed single variant. Within each evaluation we mapped in parallel simulated Illumina reads from (a) the array containing introduced variant sequence and (b) the array that lacked the variant. In experiments where reads containing the introduced variant where mapped to the reference containing the variant we observed the introduced base in across variations of sequence coverage and increased error rates. To validate a variant as “true”, we next evaluated the supporting sequence coverage. For example in 100x coverage, using the default Illumina error rate we observed 96 “true” calls out of 100 simulations, where in each case we set a threshold that at least 80% of reads that spanned the introduced variant supported the base call. We found that Illumina quality did influence our ability to confidently validate array variants by reducing the coverage. When the substitution error was increased by 1/10th we observed a decrease to only 75 "true” variant calls out of 100x simulations. Therefore, we suspect that Illumina sequencing errors may challenge our ability to completely detect true positive variants.

In our alternate experiments, whereas simulated Illumina reads from 10 identical copies of the DYZ3 repeat were mapped to a reference containing an introduced variant, we did not observe a single simulation/condition with sufficient coverage for "true” validation. We do report an increase in the percentage of reads that support the introduced variant as we increase the Illumina substitution error rate, however the range of read depth observed across all experiments were far below our coverage threshold. We obtained similar results when we repeated this simulation using sequences from the RP11-718M18 DYZ3 array.

Finally, standard quality Illumina-based polishing with pilon 21 was applied strictly to unique (that is, non-satellite DNA) sequences on the proximal p and q arms to improve final quality. Alignment of polished consensus sequences from our control BAC from Xq24 (RP11-482A22) and non-satellite DNA in the p-arm adjacent to the centromere (Yp11.2, RP11-531P03), revealed base-quality improvement to >99% identity.

#### Code availability

This study used previously published software: alignments were performed using blasr (version 1.3.1.124201) and bwa mem (0.7.12-r1044). Consensus alignments were obtained using kalign (version 2.04). Global alignments of HORs used needle (EMBOSS:6.5.7.0). Repeat characterization was performed using RepeatMasker. Satellite monomers were determined using profile hidden Markov model (HMMER3). Jellyfish (version 2.0.0) was used to characterize k-mers. Additional scripts used in preparing sequences before consensus generation are deposited in GitHub: https://github.com/khmiga/CENY.

### III. Prediction and validation of DYZ3 array

BAC ordering was determined using overlapping informative single nucleotide variants (including the nine DYZ3 6.0 kb structural variants) in addition to alignments directly to either assembled sequence on the p-arm or q-arm of the human reference assembly (GRCh38). Notably, physical mapping data were not needed in advance to guide our assembly. Rather these data were provided to evaluate our final array length predictions. Full length DYZ3 HORs (ordered 1-52) were evaluated by MSA (using kalign ^26^) between overlapping BACs, with emphasis on repeats 28-35 that define the overlap between BACs anchored to the p-arm or q-arm (Supplementary Fig 13). RPC1-11 BAC library has been previously referenced as derived from a carrier a known carrier of haplogroup R1b^30,31^. We compared our predicted DYZ3 array length with 93 R1b Y-haplogroup matched individuals by intersecting previously published DYZ3 array length estimates for 1000 genome phase 1 data ^13,14^ with donor-matched Y-haplogroup information ^32^. To investigate concordancy of our array prediction with previous physical maps of the Y-centromere we identified the positions of referenced restriction sites that directly flank the DYZ3 array in the human chromosome Y assembly (GRCh38) ^6,7,33^. It is unknown if previously published individuals are from the same population cohort as the RPC1-11 donor genome, therefore we performed similar PFGE DYZ3 array PFGE length estimates using the HuRef B-lymphoblast cell line (available from Coriell Institute as GM25430), previously characterized to be in the R1-b Y-haplogroup ^34^.

#### PFGE alpha satellite Southern

High-molecular-weight HuRef genomic DNA was resuspended in agarose plugs using 5e6 cells per 100 uL 0.75% CleanCut Agarose (CHEF Genomic DNA Plug Kits Cat #: 170-3591 BIORAD). A female lymphoblastoid cell line (GM12708) was included as a negative control. Agarose plug digests were performed overnight (8-12hrs) with 30-50U of each enzyme with matched NEB buffer. PFGE Southern experiments used ¼-½ agarose plug per lane (with an estimate of 5-10ug) in an 1% SeaKem LE Agarose gel and 0.5 X TBE. CHEF Mapper conditions were optimized to resolve 0.1-2.0 Mb DNAs: voltage 6V/cm, runtime: 26:40 hrs, in angle: 120, initial switch time: 6.75 s, final switch time: 1m33.69s, with a linear ramping factor. We used the Lambda (NEB; N0340S) and S.cerevisiae (NEB; N0345S) as markers. Methods of transfer to nylon filters, prehybridization, and chromosome specific hybridization with 32P-labeled a satellite probes have been described ^35^ Briefly, DNA was transferred to nylon membrane (Zeta Probe GT nylon membrane; CAT# 162-0196) for ~24hrs. DYZ3 probe (50 ng DNA labelled ~2 cpm/mL; amplicon product using previously published STS DYZ3 Y-A and Y-B primers ^36^) was hybridized for 16 hrs at 42C. In addition to standard wash conditions ^35^, we performed two additional stringent wash (buffer: 0.1% SDS and 0.1x SSC) steps for 10 min at 72C to remove non-specific binding. Image was recovered after 20hr exposure.

### IV. Sequence characterization of Y centromeric region

The DYZ3 HOR sequence and chromosomal location of the active centromere on the human chromosome Y is not shared among closely related great apes ^37,38^. However, previous evolutionary dating of specific transposable element subfamilies (notably, L1PA3 9.2–15.8 MYA ^39^) within the divergent satellite DNAs, as well as shared synteny of 11.9 kb of alpha satellite DNA in the chimpanzee genome Yq assembly indicate that the locus was present in the last common ancestor with chimpanzee (Supplementary Fig. 14).

Comparative genomic analysis between human and chimpanzee were performed using UCSC Genome Browser liftOver ^40^ between human (GRCh38, or hg38 chrY:10,203,170-10,214,883) and the chimpanzee genome (panTro5 chrY:15,306,523-15,356,698, with 100% span at 97.3% sequence identity). Alpha satellite and adjacent repeat in the chimpanzee genome that share limited sequence homology with human were determined used UCSC repeat table browser annotation ^41^.

The location of the centromere across primate Y-chromosomes was determined by fluorescence in situ hybridization (FISH) (Supplementary Fig. 14). Preparation of mitotic chromosomes and BAC-based probes were carried according to standard procedures ^42–44^. Primate cell lines were obtained from Coriell: *Pan paniscus* (Bonobo) AG05253; *Pan troglodytes* (Common Chimpanzee) S006006E. Male gorilla fibroblast cells were provided by Dr Stephen O’Brien (National Cancer Institute, Frederick, MD) as previously discussed ^45^. The HuRef cell line ^34^ (GM25430) was provided through collaboration with Samuel Levy. BAC DNAs were isolated from bacteria stabs obtained from CHORI BACPAC. Metaphase spreads were obtained after a 1 h 15 min colcemid/karyomax (Gibco) treatment followed by incubation in a hypotonic solution. Cells were counterstained with 4',6-diamidino-2-phenylindole (DAPI) (Vector). BAC DNA probes were labeled using Alexa flour dyes (488, green and 594, red) (ThermoFisher). The BAC probes were labeled with biotin 14-dATP by nick translation (Gibco). And the chromosomes were counterstained with DAPI. Microscopy, image acquisition, and processing were performed using standard procedures.

### V. Epigenetic mapping of centromere proteins

To evaluate similarity between the HuRef DYZ3 reference model (Genbank: GJ212193) and our RP11 BAC-assembly we determined the relative frequency of each k-mer in the array (where k=21, with a 1-bp slide taking into account both forward and reverse sequence orientation using Jellyfish) normalized by the total number of observed k-mers (as shown in Supplementary Fig 15), with pearson correlation. Enrichment across the RP11 Y assembly was determined using the log transformed relative enrichment of each 50-mer frequency relative to the frequency of that 50-mer in background control (GEO Accession: GSE45497 ID: 200045497), as previously described ^5^. If a 50mer is not observed in the ChIP background the relative frequency was determined relative to the HuRef Sanger WGS read data (AADD00000000 WGSA) ^34^. As shown in Figure 2 in main text, average enrichment values were calculated for windows size 6 kb. Additionally, CENP-A and C paired read datasets (GEO Accession: GSE60951 ID: 200060951)^46^ were merged (PEAR ^29^, standard parameters) and mapped to all alpha satellite reference models in GRCh38. Reads that mapped specifically to the DYZ3 reference model were selected to study enrichment to the HOR array. The total number of bases mapped from CENP-A and CENP-C data versus the input controls was used to determine relative enrichment. Secondly, reads that mapped specifically to the DYZ3 reference model were aligned to the DYZ3 5.7 kb in consensus (indexed in tandem to avoid edge-effects), and read depth profiles were determined. To characterize enrichment outside of the DYZ3 array CENP-A, CENP-C and Input data were mapped directly to the RP11 Y-assembly. Reads mapping to the DYZ3 array were ignored. Read alignments were only considered outside of the DYZ3 array if no mismatches, insertions, or deletions were observed to the reference and if the read could be aligned to a single location (removing any reads with mapping score of 0). Sequence depth profiles were calculated by counting the number of bases at any position and normalizing by the total number of bases in each respective dataset. Relative enrichment was obtained by taking the log transformed normalized ratio of centromere protein (A or C) to Input.

